# Phosphorylation State Dictates Bacterial Stressosome Assembly and Function

**DOI:** 10.1101/2025.03.04.640944

**Authors:** Elizabeth A. Martinez-Bond, Ivanna Lopez-Ayala, Mariya Lobanovska, Lisa Qiu, Virginia Garda, Zanlin Yu, Daniel A. Portnoy, Allison H. Williams

## Abstract

Bacterial pathogens rely on their ability to sense and respond to environmental stressors to survive and maintain virulence. The stressosome, a 1.8-megadalton nanomachine, serves as a critical sensor and regulator of the general stress response. It is composed of multiple copies of three proteins RsbR, RsbS, and the kinase RsbT which together orchestrate activation of downstream stress adaptation pathways. Using cryo-electron microscopy, we solved the atomic structure of five *Listeria monocytogenes* stressosomes, capturing structural mimics of the transition between inactive and activated states using phosphomimetic and phosphodeficient mutants. Our findings reveal that phosphorylation at specific residues T175 and T209 on RsbR, and S56 on RsbS dictates stressosome assembly, stoichiometry, and activation. Specifically, phosphorylation at T175 primes the stressosome for activation, while S56 phosphorylation destabilizes the core, triggering the release of RsbT to propagate the stress response. In contrast, phosphorylation at T209 modulates stressosome composition and appears to fine-tune the intensity of the stress response. Functional analyses reveal that phosphomimetic mutants (T209E, S56D) resist oxidative stress but lose virulence in host cell model, while phosphodeficient mutants (T175A, S56A) are stress-sensitive but retain virulence. These findings establish phosphorylation as a central regulatory switch linking structural dynamics to bacterial adaptation and pathogenesis, highlighting potential targets for antimicrobial intervention.

## Introduction

Microorganisms constantly face environmental stresses, and in bacteria, the stressosome plays a central role in initiating the stress response. The stressosome, first discovered in *Bacillus subtilis*, is among the largest bacterial nanomachines^1,2^. This 1.8-MDa nanomachine responds to environmental changes by triggering a protein partner-switching cascade that activates an alternative sigma factor, σ^B^ ^3,4^. This regulatory network governs the expression of over 300 genes essential for stress resilience, virulence, and bacterial survival^5^. While the stressosome role in bacterial stress adaptation is well studied, the molecular mechanisms underlying its activation— particularly how phosphorylation regulates its assembly and function—remain poorly understood.

In *Listeria monocytogenes* (*Lm*), the stressosome governs both environmental stress adaptation and virulence modulation, playing a crucial role in bacterial survival during infection^4,6^. Stressosome signaling is needed for bacterial persistence against host-imposed stresses, such as reactive oxygen and nitrogen species, nutrient limitation, and phagosomal challenges^2^. Activation of σ^B^ directly regulates *prfA*, the master virulence transcription factor, linking stress responses to key pathogenic processes, including host invasion, intracellular replication, and intercellular spread^6,7^.

The best-studied stressosome models are from *Bacillus* and *Listeria species*^1,2,4,6,8,9^. This supramolecular structure primarily consists of multiple copies of three proteins RsbR, RsbS, and RsbT that assemble into a virus-like capsid, an atypical pseudo-icosahedron. The *C*-terminal domains of RsbR and RsbS both feature a conserved sulfate transporter and an anti-sigma factor (STAS) domain (Supplementary Figure 1a-c)^4,6^. RsbR also has an additional non-heme globin domain at its *N*-terminus that resembles protruding turrets (Supplementary Figure 1b-c). Previous studies have shown that the turret-like structures of RsbR are likely to be involved in protein-protein interactions and environmental stress sensing^10^. The STAS domains of RsbR and RsbS interact to form the core structure of stressosome that consists of 20 dimers of RsbR and 10 dimers of RsbS (Supplementary Figure 1c). The 2:1 RsbR to RsbS ratio in the stressosome complex has been consistently observed in all native structures of Gram-positive *Bacillus* and *Listeria*^1,2,6–8,11^. Interestingly, the stressosomes in Gram-negative *Vibrio brasiliensis* and *Vibrio vulnificus* have shown core composition variability of RsbR and RsbS (1:1 and 4:2, respectively)^12,13^. In these studies, RsbR and RsbS were both cloned and expressed under native conditions and potentially represent an intermediate state in activation of the stressosome.

The current model for stressosome activation in *Listeria* suggests that, in the absence of stress, the kinase RsbT is sequestered by the RsbR/RsbS core, preventing σ^B^ activation^4^. Upon stress, the stressosome is recruited to the membrane by the small protein Prli42, initiating its activation^10^. The core proteins RsbR and RsbS contain three conserved phosphorylation sites: T175 and T209 on RsbR, and S56 on RsbS (Supplementary Figure 1). Phosphorylation of T175 and S56 by RsbT triggers dissociation of RsbT from the core, allowing it to activate downstream effectors and σ^B^, thereby initiating the transcription of the stress response genes^4,5,14,15^. Phosphorylation of T209 on RsbR dampens the stress response, fine-tuning σ^B^ activation^3,6^. RsbX, the phosphatase for RsbT, presumably resets the system by dephosphorylating RsbR and RsbS^16,17^.

Here, we present five cryo-EM structures that capture key steps in the transition from an inactive to an activated stressosome. These structures reveal how the phosphorylation of RsbR T175, T209, and RsbS S56 regulates stressosome assembly, stoichiometry, and activation. Phosphorylation of T175 prepares the stressosome for activation by altering the core’s spatial structure, while phosphorylation of T209 can regulate stress activation by decreasing the incorporation of the sensory protein RsbR into the stressosome core. Phosphorylation of S56 likely destabilizes the stressosome complex, facilitating the release of RsbT to carry out its downstream activity. Combined with functional analyses of phosphomimetic and phosphodeficient mutants, these findings underscore the pivotal role of phosphorylation in balancing bacterial stress resilience and virulence. Together, our studies provide a mechanistic framework for understanding how the stressosome enables bacteria to adapt to rapidly changing environments, offering insights that may guide the development of therapeutic strategies targeting pathogenicity.

## Results

### Cryo-EM structure determination of five *Listeria* stressosome complexes

Previous studies showed that in the absence of RsbT, the stressosome did not form homogeneous stressosome particles^6^. We hypothesized that RsbT serves not only an enzymatic role in stressosome activation but also a crucial structural role in stressosome assembly. RsbR and RsbS, the primary substrates for RsbT, together constitute the core structure of the stressosome, a large truncated icosahedral signaling complex^6,7,18^. Here we leveraged the power of cryo-EM single particle analysis to investigate the role of phosphorylation in stressosome activation and assembly. To mimic the phosphorylation state, we generated single amino acid substitutions at each conserved phosphorylation site of RsbR (T209, T175) and RsbS (S56). The quality of the assembled stressosomes was first assessed by SDS-PAGE, and the homogeneity of the stressosome was evaluated by negative stain electron microscopy (Figure 1a-e, Supplementary 2a-c and 3a-c). We determined 3D reconstructions of five stressosome-containing mutants (RsbR T209 A/E, RsbR T175 A/E, and S56A). S56D did not form homogeneous stressosome assemblies, so a 3D reconstruction was not attained, suggesting stressosome formation is destabilized when S56 of RsbS is phosphorylated (See Table 1, Supplementary Figure 3b). The five mutant structures were resolved at resolutions ranging from 3.4 Å to 4.2 Å (Figure 1a-e iii, Table 1, Supplementary Figure 4). No symmetry constraints were applied during the data reconstruction process, ensuring that the maps presented reflect the original data without imposed biases (Table 1, Figure 1a-e).

**Figure 1.**
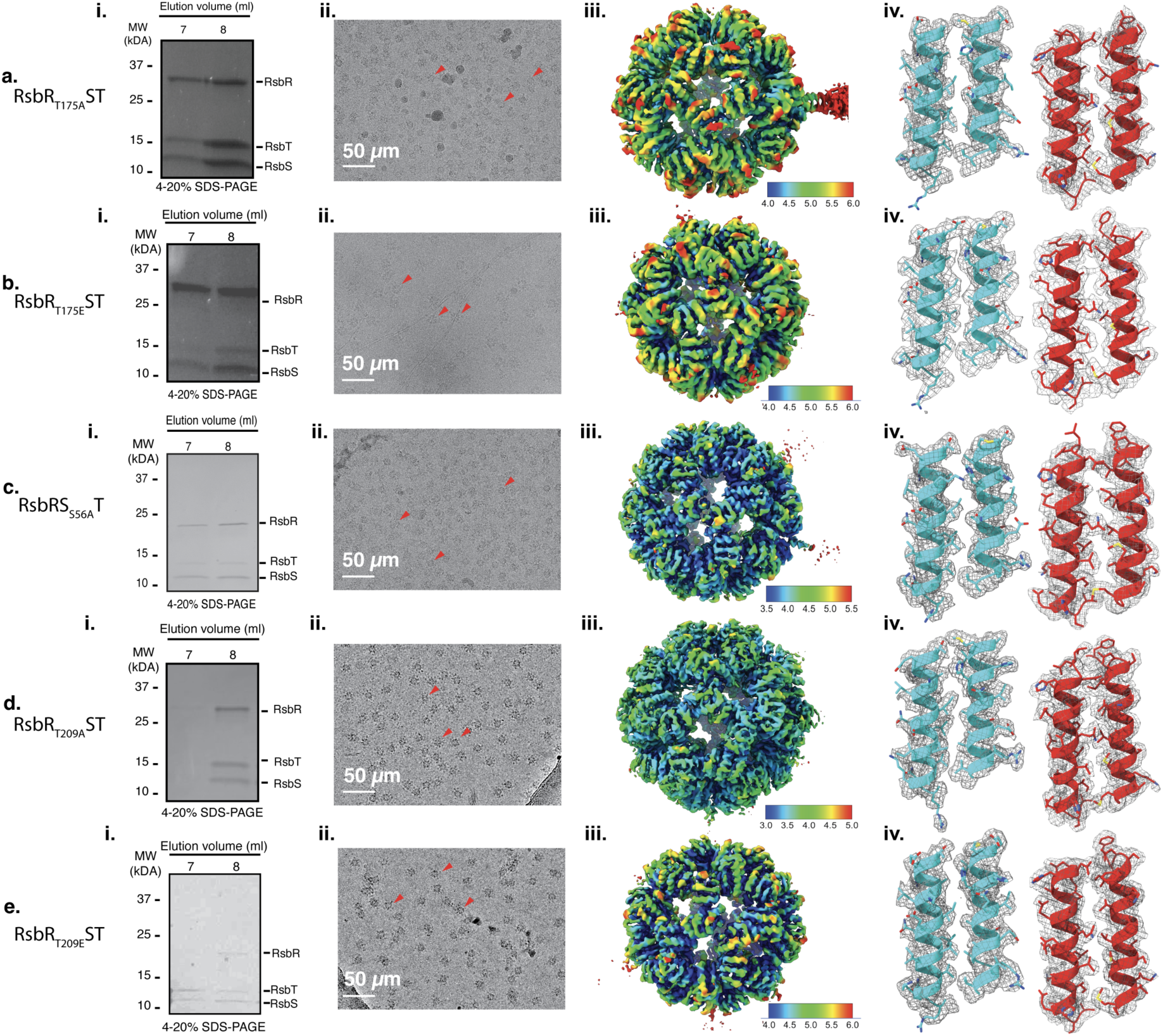
Cryo-EM structure determination of five *Listeria* stressosome complexes (a-e) Analysis of stressosome complexes for the indicated mutants: a) RsbR_T175A_ST, b) RsbR_T175E_ST, c) RsbS_S56A_RT, d) RsbR_T209A_ST, and e) RsbR_T209E_ST. *(i)* SDS-PAGE analysis of purified stressosome complexes. *(ii)* Cryo-EM micrographs of stressosome complexes, with particles highlighted by red arrows. *(iii)* Overall electron density maps of the core stressosome structures. Maps are contoured at ∼sigma level 3, with local resolution mapped according to the side color scale. *(iv)* Quality of the electron density and model fit. Left panels show RsbR, and right panels display RsbS, with helices labeled according to residue numbers 175-193 and 209-225 for RsbR and 20-41 and 53-73 for RsbS.

**Table 1.** Data Collection and Processing Summary.

| Parameter | RsbR<sub>T209E</sub>ST | RsbR<sub>T209A</sub>ST | RsbR<sub>T175E</sub>ST | RsbR<sub>T175A</sub>ST | RsbRS<sub>S56A</sub>T |
| --- | --- | --- | --- | --- | --- |
| **Data Collection and Processing** |  |  |  |  |  |
| Magnification | 105,000 | 105,000 | 105,000 | 105,000 | 105,000 |
| Voltage (kV) | 300 | 300 | 300 | 300 | 300 |
| Electron exposure (e<sup>-</sup>/Å<sup>2</sup>) | 59 | 59 | 59 | 59 | 59 |
| Defocus range (μm) | -0.8 - -2.0 | -0.8 - -2.0 | -0.8 - -2.0 | -0.8 - -2.0 | -0.8 - -2.0 |
| Pixel size (Å) | 0.8635 | 0.8635 | 0.8635 | 0.8635 | 0.8635 |
| Symmetry imposed | C1 | C1 | C1 | C1 | C1 |
| Final Particle images (no.) | 31,771 | 233,090 | 58,676 | 97,564 | 117,803 |
| Map resolution (Å) | 4.02 | 3.42 | 4.16 | 4.05 | 3.76 |
| FSC threshold | 0.143 | 0.143 | 0.143 | 0.143 | 0.143 |
| **Model Composition** |  |  |  |  |  |
| Non-hydrogen atoms | 56,610 | 58,426 | 57,860 | 57,700 | 57,760 |
| Protein residues | 7,500 | 7,724 | 7,640 | 7,640 | 7,640 |
| Ligands | - | - | - | - | - |
| B-factor (Å<sup>2</sup>) | 115.82/325.4<br>1/150.97 | 17.59/259.32<br>/84.93 | 130.91/364.1<br>5/164.81 | 47.28/365.47<br>/132.32 | 58.78/215.6<br>4/96.44 |
| **Root Mean Square (R.M.S) Deviations** |  |  |  |  |  |
| Bond length (Å) | 0.003 | 0.004 | 0.003 | 0.003 | 0.004 |
| Bond angles (°) | 0.624 | 0.725 | 0.642 | 0.744 | 0.755 |
| **Validation Metrics** |  |  |  |  |  |
| MolProbity score | 1.59 | 2.22 | 1.61 | 1.68 | 2.05 |
| Clashscore | 10.27 | 12.88 | 7.59 | 11.00 | 13.72 |
| Poor rotamers (%) | 0.06 | 3.09 | 0.03 | 0.02 | 1.76 |
| **Ramachandran Plot** |  |  |  |  |  |
| Favored (%) | 97.72 | 96.54 | 96.82 | 97.38 | 96.62 |
| Allowed (%) | 2.28 | 3.46 | 3.18 | 2.62 | 3.38 |
| Disallowed (%) | 0.00 | 0.00 | 0.00 | 0.00 | 0.00 |

### Phosphodeficient Mutants Alter Stressosome Stoichiometry and Structure in *Lm*

The archetypal stressosomes determined from Gram-positive bacteria *Listeria* or *Bacillus* have a 2:1 ratio of RsbR to RsbS in their core ^2,6,7,18^. Specifically, the stressosome core consists of 20 dimers of RsbR and 10 dimers of RsbS (Figure 2a, e1). Here we examined the role of inactivated phosphorylation sites in stressosome formation: RsbR_T175A_ST (RsbR T175A, RsbS+RsbT), RsbS_S56A_ RT (RsbS S56A, RsbR+RsbT), and RsbR_T209A_ST(RsbR T209A, RsbS+RsbT) (Figure 2b-d, e2-4). All these assemblies except for RsbR_T209A_ST maintained the precise 2:1 ratio of RsbR to RsbS in the core structure of the stressosome (Figure 2d, e4). The RsbR_T209A_ST mutant complex saw rearrangement in the stressosome core proteins, where there were 23 RsbR and 7 RsbS dimers, approximately a 3:1 ratio of RsbR to RsbS. This demonstrates the ability of RsbR T209 to determine the amount of the subunits of RsbR or RsbS incorporated into the stressosome core (Figure 2d, e4).

**Figure 2.**
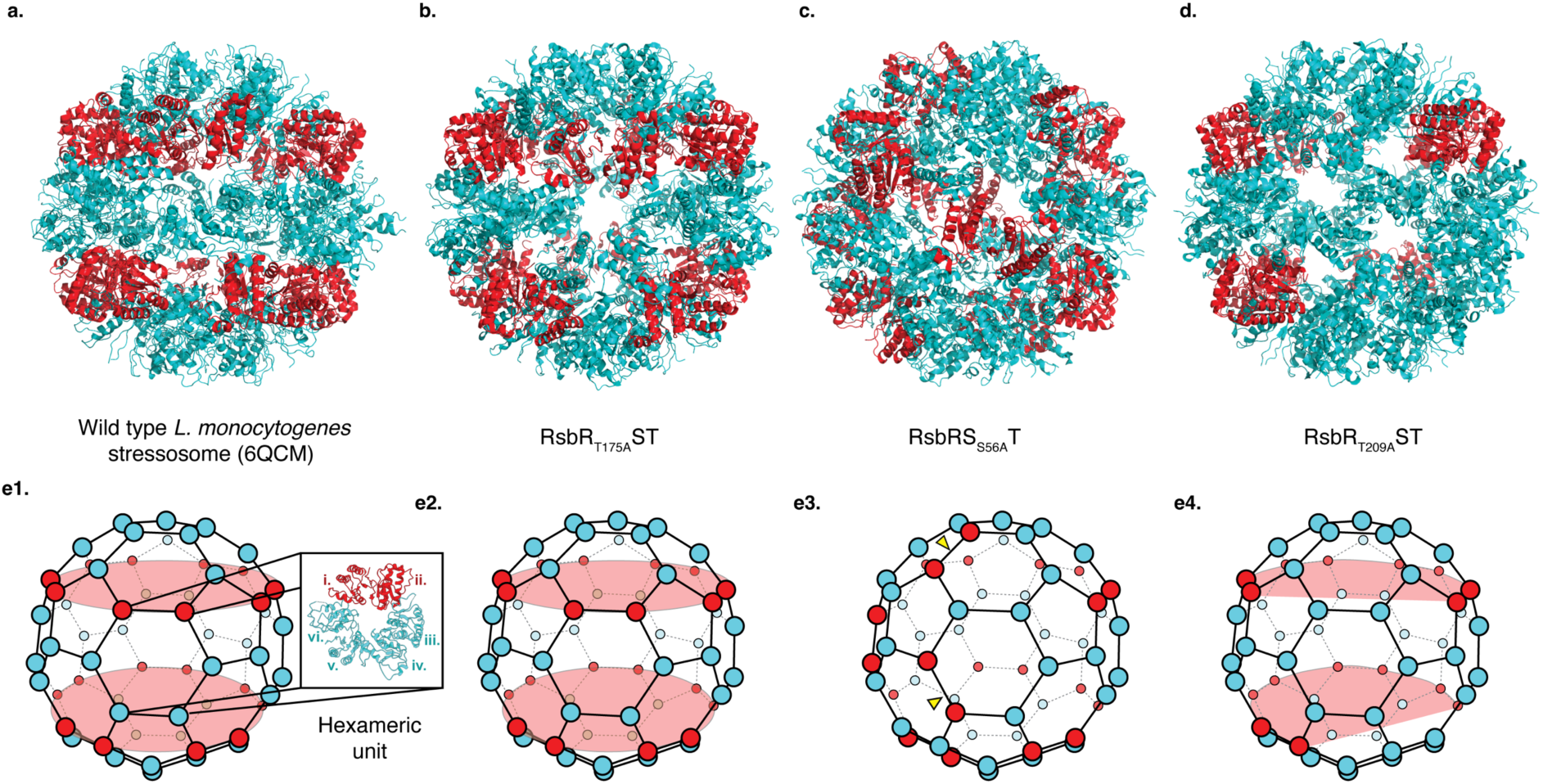
Stoichiometry of RsbR to RsbS in Inactive Stressosome Assemblies. (a) Native stressosome assembly with STAS domains of RsbR and RsbS shown in cyan blue and red ribbon representations, respectively. RsbS (red) forms two distinct bands around the stressosome core, with a stoichiometry of 20 RsbR dimers to 10 RsbS dimers. (b) RsbRT175AST mutant stressosome assembly. The overall architecture and stoichiometry are comparable to the native stressosome in Figure 2a. (c) RsbSS56ART mutant stressosome assembly. This mutant adopts a structure similar to the published *L. innocua* structure (PDB: 7B0U), where the two continuous RsbS rings are disrupted. Arrows in red point to RsbS dimers converging on a pentameric face of the stressosome. The RsbR-to-RsbS stoichiometry remains similar to that of the native complex. (d) RsbR T209AST mutant stressosome assembly. Structural rearrangements result in a stoichiometry of approximately 23 RsbR dimers to 7 RsbS dimers, reflecting a 3:1 ratio. (e1-4) Schematic representation of the stressosome assemblies 2a-d. Proteins is depicted as balls: cyan blue for RsbR and red for RsbS.

There were subtle variations in the arrangement of RsbR and RsbS in the stressosome core of RsbR_T175A_ST and RsbS_S56A_RT assemblies (Figure 2b-c, e2-3). RsbS_S56A_RT formed structures similar to the published *L. innocua* structure (7B0U), where there are no longer two continuous rings of RsbS around the core (Figure 2c, e3)^18^. Instead, there is a break in the ring where 2 RsbS dimers come together on the pentameric face of the stressosome (Figure 2c, e3). Conversely, RsbR_T175A_ST resembled *Lm* stressosome (6QCM) where there was a continuous ring of RsbS around the stressosome core (Figure 2a, e1)^10^. These results suggest that the stressosome may exist in two distinct resting states: an unprimed, native state and a primed, pre-activated state. The unprimed state resembles the wild-type stressosome and represents its native, inactive form with a highly symmetric core structure (Figure 2a-b). In the primed state, the stressosome core shows a disruption in the RsbS ring structure (Figure 2c-d). The S56A mutant represents a state where full activation of the stressosome is hindered and could represent the preactivated state of the stressosome since T175, the priming residue, remains available for phosphorylation in the RsbS_S56A_RT (Figure 2c). However, despite variations in the arrangement of RsbR and RsbS in the stressosome core among inactivated stressosome structures, the compositions of RsbR and RsbS in RsbR_T175A_ST and RsbS_S56A_RT (Figure 2b-c) are the same as the native stressosome complex (Supplementary Figure 1).

### Phosphodeficient Stressosome Mutants Are Oxidative-Stress Sensitive but Retain Virulence

To assess whether the phosphorylation state of the RsbR and RsbS components of the stressosome influences infection, we constructed phosphomimetic and phosphodeficient mutants in *Lm* at the native stressosome locus. We then evaluated their growth in rich medium (BHI) and defined medium (iLSM) supplemented with the reducing agent TCEP. The combination of iLSM and TCEP has been shown to activate the *Lm* virulence program and mimic the intracellular reducing environment encountered during infection ^19,20^. All the mutants displayed growth phenotypes comparable to WT and Δstressosome in both BHI and iLSM + TCEP (Figure 3a-b), suggesting that alterations in stressosome phosphorylation state do not affect bacterial growth.

**Figure 3.**
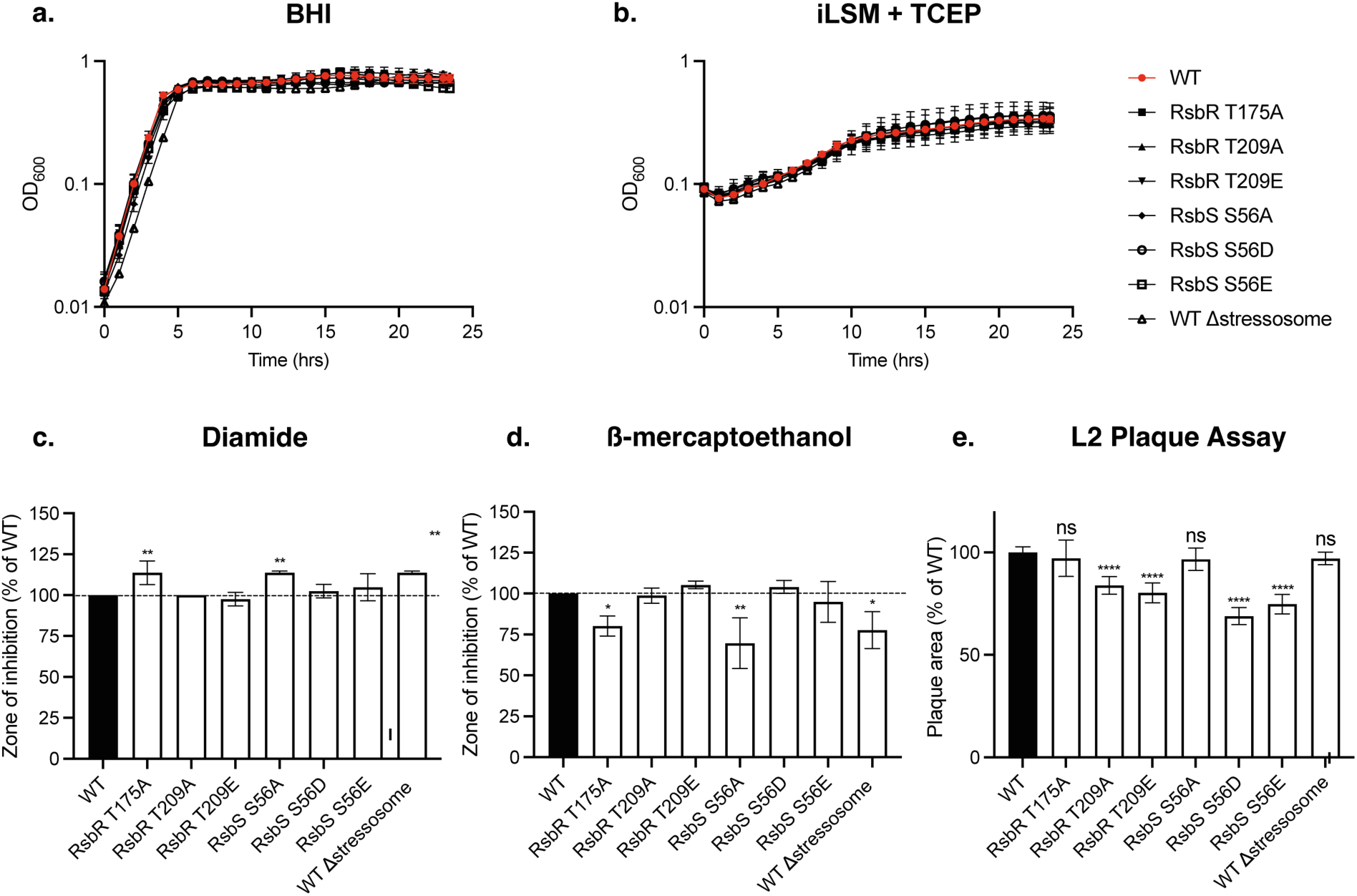
Stressosome mutants respond differently to host-mimicking stress signals and impact *Lm* virulence in cells. Growth of the indicated mutants in BHI broth culture (a) and in defined culture media iLSM supplemented with a chemical reducing agent TCEP (b). Overnight cultures were grown in BHI and then diluted in fresh media. OD600 was measured every hour for 24h. Error bars indicate standard deviation. The data represents three independent biological replicates.(c-d) Phosphorylation mutants show difference in sensitivity to redox stressors including (c) diamide and (d) b-mercaprtoethanol. One-way ANOVA with multiple comparisons to WT was used. *P <0.05; **P < 0.01. (e) Mutants with different phosphorylation state display virulence defects in fibroblasts plaque assay. One-way ANOVA with multiple comparisons to WT was used to calculate p value. ns, not significant, ****P < 0.0001.

To test whether the stressosome phosphodeficient mutants can respond to host-mimicking stress signals, we tested the mutants in their ability to survive oxidative stress and reductive stress. We used thiol-specific oxidant diamide to mimic reactive oxygen species as well as reactive electrophilic species stress response^21^. Exposure to diamide results in formation of disulfide bonds between proteins and low molecular thiols such as glutathione, which is essential for *Lm* virulence^22,23^. RsbR and RsbS mutants that have impaired stressosome signalling (RsbR T175A and RsbS S56A) were more sensitive to diamide using disc diffusion assay (Figure 3c), suggesting that the functional stressosome signaling is necessary for bacterial survival during oxidative stress. Next, we tested survival of the mutants in response to thiol reductant beta-mercaptoethanol (β-ME). Thiol reductive stress response has previously been shown to be important during *Mycobacterium tuberculosis* infection and is mediated by a number of sigma factors including σ^B^ ^24^. In contrast to the response to diamide, *Lm* stressosome mutants that had inactive stressosome signaling core (RsbR T175A and RsbS S56A) were more resistant to β-ME, similar to Δstressosome (Figure 3d). This implies that turning off the stressosome signalling can increase bacterial fitness under reductive stress.

Next, we examined the virulence of the mutants in L2 fibroblast infection assay (Figure 3e). Following a three-day infection, bacterial plaque formation serves as a measure of intracellular growth and cell-to-cell spread. RsbS S56A and RsbR T175A mutants in *Lm* displayed no defects in invasion, effectively infecting host cells and generating plaques of similar size to the wild type (Figure 3e). Similarly, Δstressosome exhibited no plaque defect suggesting that stressosome activity is not essential for virulence in L2 plaque infection model. In contrast, RsbR T209A mutant, containing a rearranged ratio of the core stressosome proteins, displayed virulence defect (Figure 3e). This suggests that proper stressosome signalling regulation at RsbR T209 is necessary for full virulence of *Lm* during infection.

### Phosphomimetic Mutations Alter Stressosome Stoichiometry, Structure, and Activation

The core proteins of the stressosome (RsbR, RsbS, and RsbT) undergo phosphorylation, triggering downstream signaling. However, the structural changes that enable phosphorylation and how these transitions propagate within the complex remain elusive. We are interested in specific conformational changes that occur when the stressosome is activated. In our previous studies, we showed that there is a flexible loop that could block access to T209 suggesting the position of the loop could be determined by the activating kinase RsbT^6^. Furthermore, we hypothesized above that the stressosome could have two distinct resting states: an unprimed, native state and a primed, pre-activated state. To address these questions, we used three phosphomimetic assemblies: 1) RsbR_T175E_ST(RsbR T175E, RsbS+RsbT), 2) RsbS_S56D_RT (RsbS S56D, RsbR+ RsbT), and 3) RsbR_T209E_ST (RsbR T209E, RsbS+RsbT).

The structure of RsbR_T175E_ST showed no changes in stoichiometry of the core stressosome components. However, similar to the published *L. innocua* structure (7B0U) and RsbS_S56A_RT described above, there are no longer two continuous rings of RsbS around the core (Figure 2c, e3, Figure 4a, c1). This suggests that the stressosome is primed and ready for activation, leading to structural readjustments in the core structure to prepare for phosphorylation of the activating residue S56. Conversely, the RsbR_T209E_ST mutant altered the composition of the complex, reducing RsbR and increasing RsbS subunits, resulting in a 1:1 ratio of RsbR to RsbS with 15 dimers of each protein (Figure 4b, c2). This structure is the only present example in Gram-positive bacteria where the number of RsbR molecules decreased in the core of the stressosome. These findings align with the mechanism where T209 phosphorylation decreases stress activation by limiting RsbR integration into the stressosome (Figure 4b, c2).

**Figure 4.**
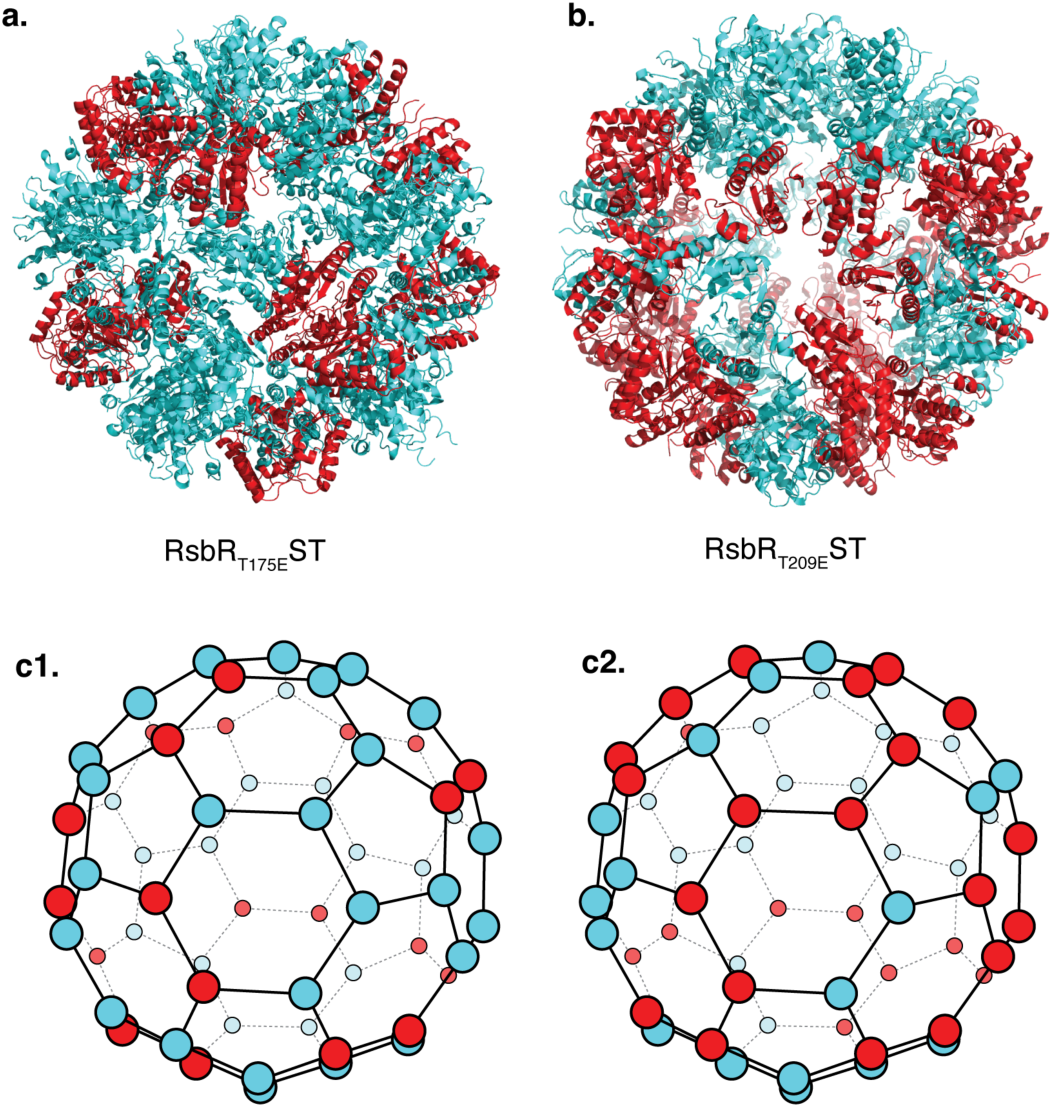
Structures Mimicking Activated Stressosome Complexes. (a) RsbRT175EST mutant stressosome assembly. The architecture is similar to RsbSS56AST structure shown in Fig. 2c, maintaining a stoichiometry of 20 RsbR dimers and 10 RsbS dimers, comparable to the native stressosome. (b) RsbRT209EST mutant stressosome assembly. This mutant exhibits a 50% reduction in RsbR within the complex, resulting in a stoichiometry of 15 RsbR dimers and 15 RsbS dimers.

The evaluation of RsbS_S56D_RT mutant by negative stain electron microscopy repeatedly revealed an absence of homogenous formation of the stressosome. Instead, stressosome particles were dispersed within a field of debris, consisting of partial or broken stressosome particles (Supplementary Figure 3b). Since a high-resolution 3D reconstruction by cryo EM of RsbS_S56D_RT was not feasible, it is plausible that the stressosome could be destabilized when this residue is phosphorylated, and this is how RsbT is released to perform its downstream activities.

### T209 Phosphomimetic Mutation Modulates RsbR Integration into Stressosome Core

Our studies reveal that the stressosome is a far more dynamic structure than previously understood. In its native state, the basic unit of the stressosome complex forms a heterotrimer consisting of two RsbR homodimers and one RsbS homodimer (Supplementary Figure 5a-d)^6^. However, the RsbR_T209E_ST mutant induces a significant alteration in stoichiometry between RsbR and RsbS in the stressosome core (Figure 4b,c2). This mutant produces a unique variant of the stressosome that not only retains the native heterotrimeric assembly within the core (Figure 5 Unit 1, a-d), but also forms a novel heterotrimer, consisting of one RsbR dimer flanked by two RsbS dimers (Figure 5 Unit 2, e-h).

**Figure 5.**
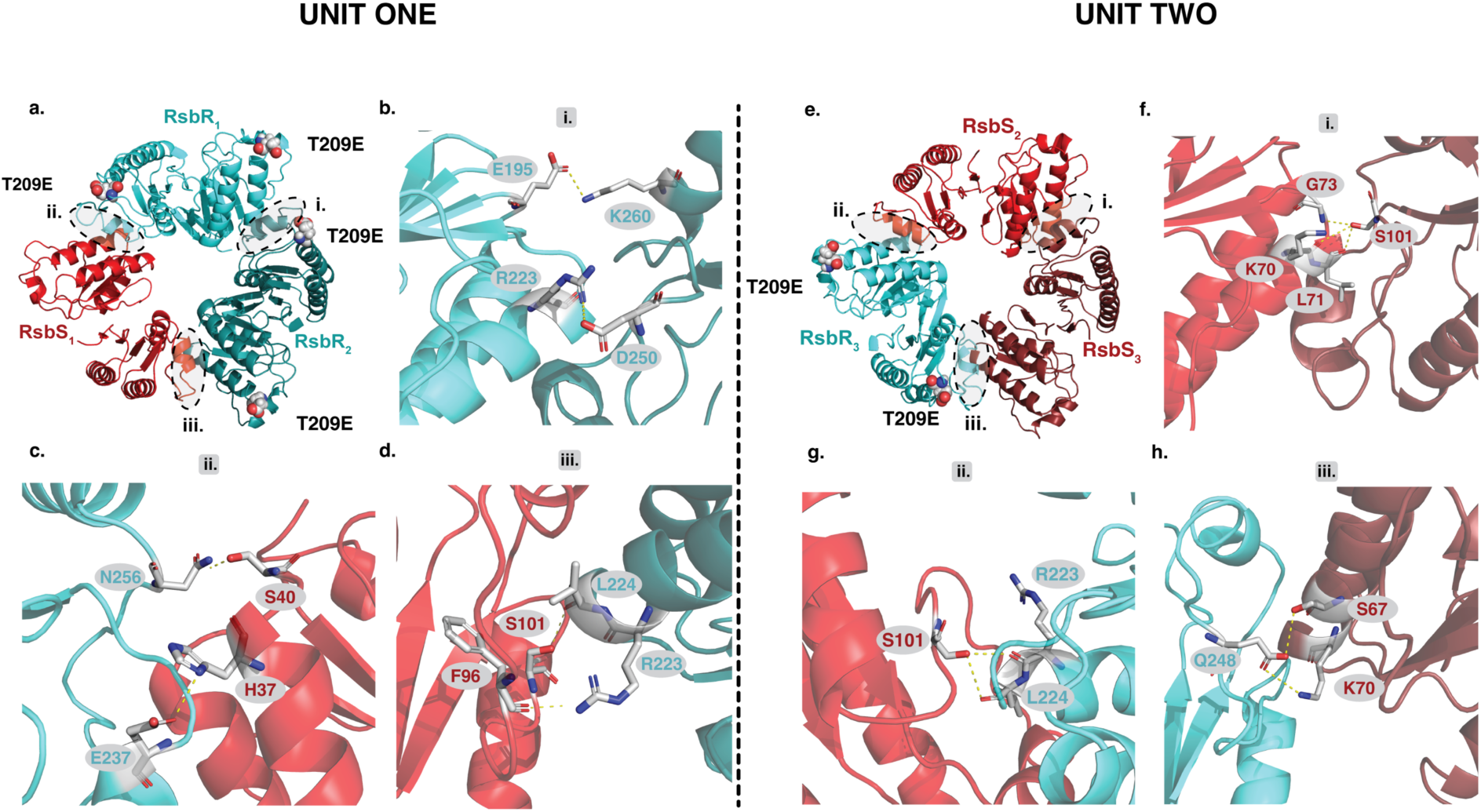
The Hexameric Units of the Mutant RsbRT209E ST Complex. a) Typical heterotrimeric assembly of RsbRT**209E**ST. Unit one is similar to the native stressosome complex. It consists of a heterotrimer composed of two RsbR homodimers and one RsbS homodimer. This unit forms three distinct interfaces: (i) RsbR–RsbR, (ii) RsbR–RsbS, and (iii) RsbR–RsbS. Each unit contains four T209E residues: two located within Interface i, facing each other, and one in each of Interfaces ii and iii. (b-d) Interfaces i, ii and iii are primarily held together by salt bridge and hydrogen bonds. See supplementary table 1 for a detail description of all the interactions. e) Unique heterotrimeric assembly of the stressosome variant RsbRT**209E**ST. Unit two stressosome variant forms a distinct heterotrimeric assembly. The new heterotrimer consists of one RsbR dimer flanked by two RsbS dimers, held together by three interfaces: (i) RsbS–RsbS, (ii) RsbS–RsbR, and (iii) RsbS–RsbR. The assembly includes two T209E residues, located at Interfaces ii and iii, representing a reduction of two T209 residues compared to the native heterotrimer arrangement in Figure 4a and Supplementary Figure 6. f-h) Interfaces i, ii and iii are held together by salt bridges and hydrogen bonds. See supplementary table 1 for detailed description of all the interactions.

Unit 1 of RsbR_T209E_ST stressosome has three interfaces: one between two RsbR dimers (Figure 5a, Interface i) and two between RsbR and RsbS dimers (Figure 5a, Interfaces ii and iii). These interactions are stabilized by hydrogen bonds and salt bridges (Figure 5b-d, Supplementary Table 1). In this configuration, four E209 residues are present: two at interface i, facing each other, and one each at interfaces ii and iii (Figure 5a).

In Unit 2, the RsbR_T209E_ST stressosome maintains all three interfaces but features a rearranged architecture. Specifically, it includes two interfaces between RsbR and RsbS dimers (Figure 5e, Interfaces ii and iii) and one interface between two RsbS dimers (Figure 5e, Interface i). This altered arrangement reduces the number of E209 residues to two, located at interfaces ii and iii (Figure 5e), compared to the four E209 residues in the native heterotrimer (Figure 5a, Supplementary Figure 5a^6^). Like the native complex, the interfaces in the mutant are stabilized by hydrogen bonding and salt bridges (Figure 5f-h, Supplementary Table 1).

The RsbR_T209E_ST mutant exhibits substantially fewer residues involved in protein-protein interactions than the native stressosome structure, suggesting that the mutant adopts a more dynamic and transient interaction network (Figure 5, Supplementary Figure 5a-d). The T209E mutation, which mimics phosphorylation at this position, likely serves as a regulatory mechanism modulating RsbR within the stressosome core. Interestingly, as previously mentioned, the phosphorylation-equivalent residue in *Bacillus* T205 (T209) has been associated with a diminished stress response, further reinforcing this hypothesis^3,6^.

### Phosphomimetic Stressosome Mutants Survive Oxidative Stress but Exhibit Reduced Virulence

The phosphomimetic mutations T209E in RsbR and S56D in RsbS reveal distinct roles in modulating stressosome-mediated stress responses in *Lm*. In our disk diffusion assay, described above Δstressosome was more sensitive to oxidative stress (diamide). In contrast, T209E and the S56 phosphomimetic mutants, S56D and S56E, exhibited a zone of inhibition comparable to that of the wild-type strain. Since WT stressosome response to diamide phenocopied phosphomimetic mutants, we conclude that WT stressosomes are activated and the phosphorylated residues are required for optimal oxidative stress resistance (Figure 3c). In contrast, the S56 phosphomimetic mutants, S56D and S56E, exhibited a zone of inhibition comparable to that of the wild-type strain, suggesting that phosphorylation at this residue does not significantly alter oxidative stress resistance (Figure 3c). Under reductive stress (β-ME), T209E, S56D, and S56E mutants displayed increased susceptibility, indicating that these modifications do not confer protection in a reducing environment (Figure 3d). Interestingly Δstressosome shows increased resistance to reductive stress (β-ME) compared to wild-type indicating that there may be a fitness cost associated with stressosome activation. Consistent with this, T209E, S56D, and S56E phosphomimetic mutants behaved the same as WT (Figure 3d). These findings suggest that while T209 and S56D/E phosphorylation is necessary for robust stressosome activity under oxidative stress, these modifications do not confer protection in a reducing environment. Despite multiple attempts, we were unable to generate an RsbR T175E mutant, suggesting that this mutation may be detrimental to bacterial viability.

We then assessed the impact of these phosphomimetic mutations on the virulence of *Lm* in L2 fibroblast infection assay. All phosphomimetic mutants (T209E, S56D, and S56E) displayed significantly impaired plaque formation compared to the wild-type (WT) strain, suggesting potential virulence defects (Figure 3e). These defects imply that the phosphomimetic substitutions may disrupt the phosphorylation-dephosphorylation cycle critical for stressosome activation, potentially affecting bacterial signaling pathway *in vivo*.

## Discussion

Our study reveals the structural and functional roles of phosphorylation in *Lm* stressosome assembly and activation, highlighting the interplay between stress response and virulence. Specifically, we show the RsbT-mediated phosphorylation of key residues—RsbR T175, RsbR T209, and RsbS S56—modulates both stressosome stoichiometry and subunit arrangement. Phosphorylation of these key residues drives transitions between inactive, primed, and active states (Figures 2, 4, and 6). These findings align with our previous work, which demonstrated the dual role of RsbT in catalyzing and stabilizing stressosome assembly, while also illustrating how phosphorylation state alterations influence stressosome integrity and resilience^6^.

Our cryo-EM analyses show that stressosome integrity is highly sensitive to phosphorylation at conserved residues. Phosphomimetic mutations disrupt core assembly (T175E and T209E), and produce heterogeneous stressosome structures (S56D), indicating instability (Figure 4a-b, Supplementary Figure 3b). In contrast, the phosphodeficient variants (T175A and S56A) retained the canonical 2:1 RsbR to RsbS stoichiometry, demonstrating the relative structural stability of unphosphorylated stressosomes (Figure 2a-c). RsbR_T209A_ST deviated from this pattern, exhibiting a noncanonical RsbR to RsbS (23:7) stoichiometry, indicating that T209 may act as a regulatory hub, dictating subunit composition within the stressosome core (Figure 2d, e4). T175 serves as the priming residue, initiating stressosome activation. Our structural data suggest that the T175E phosphomimetic locks the stressosome in a preactivated state (Figure 4a, Figure 6). The inability to generate a RsbR T175E mutant in *Lm* implies that this substitution may induce constitutive activation, resulting in persistent stress signaling and bacterial toxicity. We hypothesize that continuous RsbT release from the stressosome could overwhelm stress-response pathways, disrupt cellular homeostasis, and impair essential downstream signaling networks, ultimately leading to lethality.

**Summary Figure 6.**
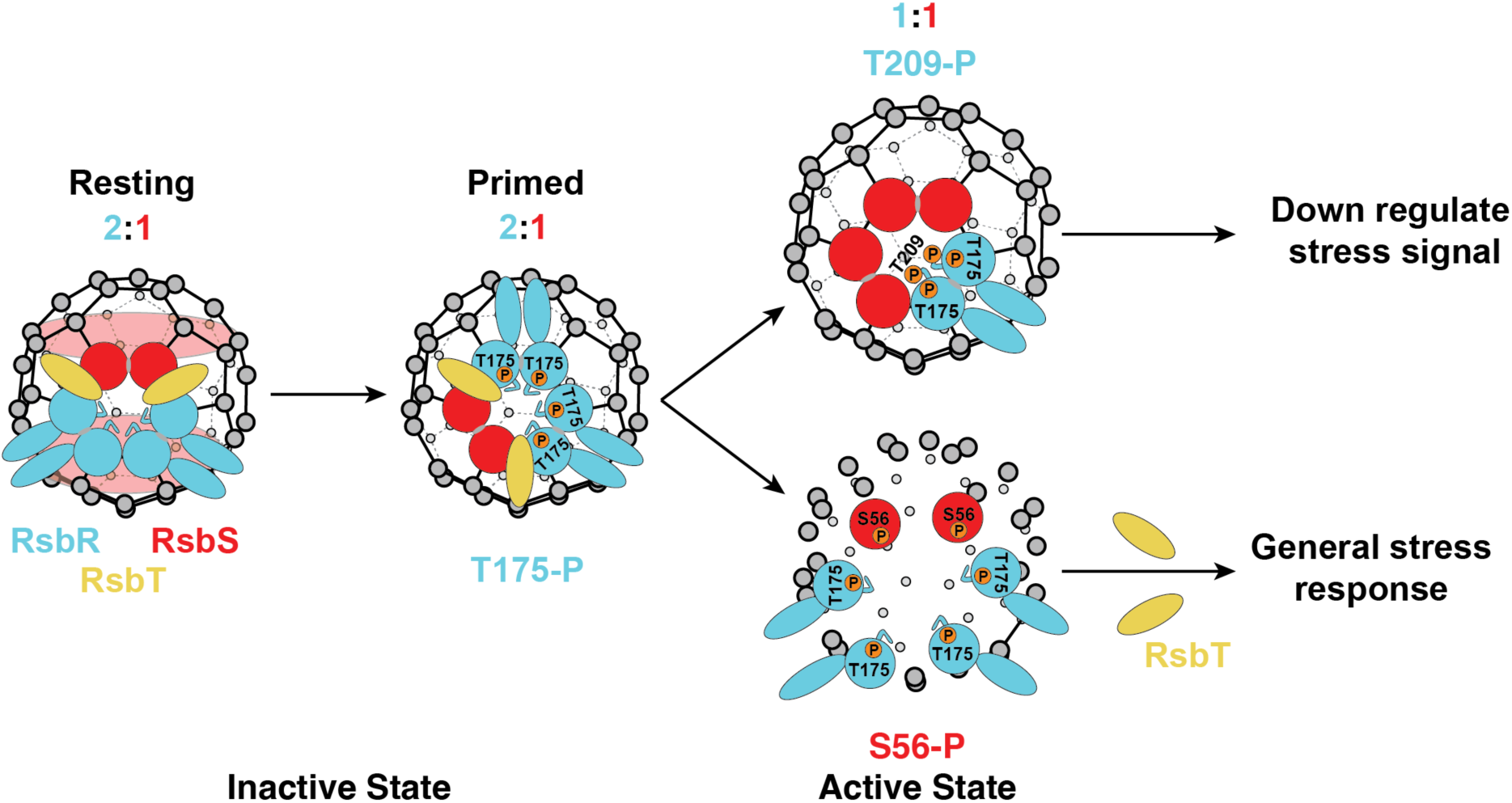
**Stressosome Activation Mechanism and Signal Integration. Inactive State**: In the resting state, the stressosome adopts a stable icosahedral-like capsid structure composed of a 2:1 ratio of RsbR (sensor) to RsbS (scaffold). This configuration sequesters the kinase RsbT, effectively preventing downstream signaling. In the primed state, upon encountering environmental stress, such as oxidative or acidic conditions, the globin-like turrets of RsbR detect these signals. In response, RsbT phosphorylates RsbR at T175, priming the stressosome. This phosphorylation disrupts the RsbS ring, preparing the structure for activation. **Active State**: Phosphorylation of RsbS destabilizes the stressosome core, leading to the release of RsbT. Freed RsbT then phosphorylates downstream effectors, activating σB. This initiates the transcription of stress response genes. **Stress Signal Regulation**: Phosphorylation at RsbR T209 modulates the response intensity by reducing the amount of sensory protein RsbR within the stressosome core, fine-tuning the stress response.

Our functional analyses highlight distinct functions for RsbR T175, T209 and RsbS S56 phosphorylation in stress adaptation. During the intracellular infection *Lm* is exposed to a range of stress signals^22^. For example, reactive oxygen species are encountered during the phagosomal stages of infection whereas in the cytosol bacteria are experiencing a reducing environment^25^.We show that the stressosome activation *via* phosphomimetic mutations is important for resistance to oxidative stress (Figure 3c), suggesting that the stressosome might be necessary during early phagosomal stages of infection. On the other hand, the downregulation of the stressosome activity through phosphodeficient T175A and S56A mutations renders bacteria more resistant to reductive stress (Figure 3d). It is possible that once in the cytosol, bacteria may actively downregulate the stressosome signaling to promote virulence and cell to cell spread in response to reducing environment. Indeed, in L2 infection model the RsbR T175A and RsbS S56A mutants, which are unable to transduce the stress signal, exhibited no virulence defect, similar to Δstressosome, whereas phosphomimetic mutants were attenuated (Figure 3e). Interestingly T209A mutant that shows non canonical rearrangement of RsbR and RsbS (Figure 2d, e4) behaved similarly to phosphomimetic mutants in both infection and in stress sensitivity assays (Figure 3c-e), consistent with our hypothesis that T209 may act as a regulatory hub.

The defect observed with phosphorylation mutants in our cell based model of infection and in disc diffusion assay was not due to difference in growth as all the mutants displayed similar growth pattern when tested in broth (Figure 3a,b). Together these data further support the model that *Lm* may downregulate the stressosome activity to optimize survival and dissemination. In line with this observation, RsbX phosphatase mutant which is defective in the ability to dephosphorylate RsbR and RsbS and therefore would presumably mimic the phosphomimetic mutants, was similarly attenuated in L2 infection model^26^. In the cytosol, *Lm* induces the expression of the virulence factor ActA, which is essential for host actin polymerization and cell-to-cell spread. Interestingly, *rsbX* mutant displayed lower ActA levels during *Lm* macrophage infection^26^. We speculate that disrupted phosphorylation-dephosphorylation cycles in phosphomimetic mutants may similarly result in lower ActA levels during infection that would account for the virulence defect in these mutants (Figure 3e).

σ^B^ has been shown to directly regulate the expression of the master virulence regulator *prfA* that is essential for virulence, suggesting a role of stressosome-dependent stress response in virulence^27–29^. However, deletion of σ^B^ results in variable phenotypes depending on the infection model; it is dispensable in an intravenous mouse model^30^ but essential for virulence in a guinea pig model^31^. Our infection model allowed us to dissect the physiological relevance of stressosome phosphorylation dynamics in *Lm* pathogenesis. We show that the absence of functional regulation of the key phosphorylation residues of RsbR and RsbS results in virulence attenuation.

Our results provide a mechanistic framework for stressosome activation. In its resting state, the stressosome forms a stable icosahedral-like capsid with a 2:1 stoichiometry of RsbR (sensor) to RsbS (scaffold), sequestering the kinase RsbT to prevent downstream signaling (Figure 6). Environmental stress signals, such as oxidative or reductive stress, are likely detected by RsbR’s globin-like turrets^18,32^. RsbT phosphorylates RsbR at T175, priming the structure and disrupting the RsbS ring, giving a less ordered stressosome. This is followed by the phosphorylation of RsbS at S56, which further destabilizes the stressosome core and releases RsbT, which transitions the stressosome into an activated state (Figure 6). RsbT then phosphorylates downstream effectors, leading to activation of σ^B^, which drives transcription of stress response genes such as *prf*A^1,8,33^. Phosphorylation at RsbR T209 modulates the intensity of the response, tailoring survival to specific stress types.

Finally, the stressosome’s ability to shift stoichiometry and undergo structural rearrangements reflects its adaptability to various stress conditions and signal transduction capabilities. Phosphorylation at T175, S56, and T209 orchestrates transition between inactive, primed, and activated states; regulating the stress response intensity and specificity. These phosphorylation dynamics link environmental resilience to *Listeria* pathogenesis, as T209-dependent modulation of stressosome activity is vital for host invasion and intracellular survival.

Together, our findings provide new insights into the molecular basis of Lm virulence, highlighting the stressosome as a central regulatory hub that integrates environmental cues with bacterial survival and pathogenesis. Beyond its role in bacterial stress signaling, the stepwise phosphorylation-driven conformational cycling of the stressosome resembles regulatory mechanisms seen in eukaryotic signalosomes, such as the inflammasome^34–36^ and Cullin-RING ubiquitin ligases^37^.This suggests that dynamic, phosphorylation-mediated multiprotein complex assembly may be a conserved principle across cellular signaling networks, with implications extending beyond microbiology into broader areas of structural and cell biology.

## Methods and Materials

### Stressosome construct cloning, protein expression and purification

#### Construct cloning

The genes encoding the wild-type proteins *rsbR*, *rsbS*, and *rsbT* were synthesized by GenScript and cloned into the pGEX-4T1 vector. Point-mutant constructs (T209E, T209A, T175E, T175A, S56A, S56D) were then generated using site-directed mutagenesis with the Q5® Site-Directed Mutagenesis Kit (New England Biolabs) (Supplementary Table 2). *Protein expression and purification* The GST-tagged RsbR, RsbS, RsbT, and their mutants were expressed in BL21 Gold (DE) competent cells (Agilent). Cultures were grown in LB media at 37 °C with shaking at 170 rpm. Induction was performed with 1 mM IPTG when the optical density at 600 nm (OD600) reached 0.6–0.8. Cells were collected after 16 hours of induction at 16 °C by centrifugation at 4000 x *g* for 40 minutes at 4 °C. Pelleted cells were resuspended in 50 mM Tris (pH 8.5), 150 mM NaCl, 1 mM DTT, and one EDTA-free protease inhibitor cocktail tablet (Roche). The resuspension was sonicated at 4 °C and then ultracentrifuged at 40,000 x *g* for 40 minutes at 4 °C to obtain a clear lysate free of cell debris. GST-tagged proteins were recovered using GST Agarose Beads (Thermo Fisher Scientific). The GST tag was cleaved using thrombin (Sigma-Aldrich) at a ratio of 1:4. The individual proteins were purified using size-exclusion chromatography, selecting either Superdex200 Increase 10/300 GL or Superdex75 Increase 10/300 GL (Cytiva) in a buffer of 20 mM Tris-HCl (pH 8.5), 250 mM NaCl, and 1 mM DTT. For RsbR and RsbS, as well as their mutants, ion-exchange chromatography was conducted with a Resource Q anion exchange column (Cytiva) under the same buffer conditions, employing a NaCl gradient from 50 mM to 1000 mM. RsbT was purified using Resource S cation exchange chromatography at 20 mM Tris-HCl (pH 7.0), 1 mM DTT, with a similar NaCl gradient. Protein quality and purity were assessed using SDS-PAGE electrophoresis on Mini-PROTEAN® TGX Precast Gels with a 4-20% polyacrylamide gradient (Bio-Rad). Purified proteins were stored at 4 °C for short-term use or at-80 °C for long-term storage and preparation of protein assemblies.

### Stressosome assembly

Purified RsbR, RsbS, and RsbT proteins were combined at a 1:2:5 molar ratio and gently mixed by inversion overnight at 4 °C to facilitate complex assembly. The mixture was then loaded onto a Superose 6 Increase 10/300 GL column pre-equilibrated with buffer containing 20 mM Tris-HCl (pH 8.5), 250 mM NaCl, 1 mM DTT, and 1 mM adenosine 5’diphosphate sodium salt. The assembled protein complexes were separated from unbound components by size-exclusion chromatography, and the composition of each eluted fraction was analyzed by SDS-PAGE.

### Negative stain, Cryo-EM grid preparation, and data acquisition

For negative staining, 3 µL of purified stressosome protein complex was applied to glow-discharged 400-mesh continuous carbon grids and stained with 1% (w/v) uranyl acetate. Grids were imaged on a Tecnai T12 microscope (Thermo Fisher Scientific) operating at 120 kV, equipped with a Rio camera (Gatan Inc.). Images were recorded at a magnification of 26,000× with a defocus of ±1.0 µm.

For single-particle cryo-electron microscopy (cryo-EM), gold Quantifoil R1.2/1.3 200-or 300-mesh grids (Ted Pella Inc.) were coated with graphene oxide and functionalized on the carbon side with ethylenediamine (GO-amino). Purified stressosome protein complexes (50–150 µg/mL) were applied to the GO-amino grids (3 µL), blotted using Whatman No. 1 filter paper, and plunge-frozen in liquid ethane using a Mark IV Vitrobot. Freezing conditions included a wait time of 30 s, a blot force of 0, and blotting times ranging from 4 to 10 s at room temperature, and >90% humidity.

Cryo-EM grids were screened on either a Talos Arctica or Glacios microscope before data collection. High-resolution datasets were acquired on a Talos Arctica or Titan Krios microscope (Thermo Fisher Scientific) operating at 200 kV or 300 kV, respectively. The Titan Krios was equipped with a BioQuantum post-column energy filter and a K3 direct electron detector (Gatan Inc.). Movies were recorded automatically using SerialEM at a magnification of 105,000×, with a total electron dose of 60 e⁻/Å² and a defocus range of-1.5 to-2.0 µm. Data collection statistics are summarized in Table 1.

### Image processing

During on-the-fly image acquisition, dose-weighted micrographs were motion-corrected using MotionCor2 ^38^ via the Scipion framework. Corrected micrographs were then imported into cryoSPARC^39^ (v4.2.1) for downstream processing. Contrast transfer function (CTF) estimation was performed using CTFFIND4^40^ or Patch CTF in cryoSPARC^39^.

Particles were automatically picked using the blob picker tool in cryoSPARC, extracted, and subjected to 2D classification. Initial 3D models were generated using the ab-initio reconstruction module in cryoSPARC. The resulting density maps were subsequently refined through iterative rounds of homogenous or heterogeneous refinement to enhance resolution and quality.

### Model building and refinement

Stressosome structures were built by manually fitting AlphaFold-predicted models of RsbR and RsbS into cryo-EM density maps. These initial models were generated and positioned using UCSF ChimeraX^41^. Models were then iteratively built and refined using Coot for manual adjustments and real-space refinement in PHENIX^42^.

Wherever possible, previously unresolved density corresponding to portions of the RsbR N-terminal domain was modeled into the map. Final structure visualization and representations were created using UCSF ChimeraX^41^ or Pymol^43^.

### Broth Growth Curve

Broth growth curve assays were performed in brain heart infusion (BHI; BD bioscience) or in defined media iLSM supplemented with TCE^19,20^. Overnight cultures were grown in BHI at 37 °C shaking. The cultures were normalised to an OD_600_=0.03 in fresh BHI and to an OD_600_=0.15 in iLSM + 2mM TCEP and 200 μl of the corresponding cultures were transferred to a clear flat bottom 96 well plate (Greiner Bio). Bacteria were grown at 37°C with agitation every 15 min in a Tecan Spark plate reader. The absorbance (OD_600_) was measured every 15 min. The experiment was performed three times with each biological repeat done in technical duplicates. The growth curves show hourly OD_600_ measurements.

### Redox Stress Disc Diffusion Assay

*Lm* was grown overnight in BHI at 30°C shaking. 10^2^ overnight culture were mixed with 4ml of top agar (55°C, 0.8% NaCl and 0.8% agarose) and spread over BHI plates. 5 μl of 1 M diamide solution (Sigma) or 7M of β-mercaprtoethanol (Sigma) were added to sterile filter paper discs and the discs were placed in the center of the agar plates containing the bacterial top agar overlay. The plates were incubated at 37°C overnight and the diameter of the zone of inhibition was measured.

### L2 plaque assay

L2 fibroblasts were grown in DMEM medium supplemented with 10% FBS, 2 mM L-glutamine and 1mM of sodium pyruvate. A six well dish was seeded with 1.2 x 10^6^ L2 cells per well and incubated overnight at 37°C. *Lm* grown overnight at 30°C and L2 cells were infected at the MOI of 300. Following 1 hour infection, the cells were washed twice with PBS and 3ml of DMEM plus 0.7% agarose and 10 μg/ml gentamicin was added to each well. 48 hours post infection the plaques were stained with 2 ml DMEM containing 0.7% agarose, 10 μg/ml gentamicin and 25 μl/ml neutral red dye (Sigma N62664). 72 hours post infection the plaques were imaged, and the plaque area was quantified using ImageJ software. The experiment was performed three times for each mutant strain.

### Bacterial culture strains and genetic manipulations

*E. coli* and *Lm* strains used in the study are listed in Supplementary Table 3. *E. coli* strains were grown in LB and *Lm* strains were grown in BHI unless otherwise stated. The following antibiotics were used for selection: for *Lm* streptomycin at 200 µg/ml and chloramphenicol at 7.5 µg/ml and for *E. coli* carbenicillin at 100 µg/ml. *Lm* harbouring different *rsbR* and *rsbS* alleles were constructed by allelic exchange using pKSV7-oriT as described previously^44^. Primers used for construction of *Lm* strains are listed in Supplementary Table 2. The resulting mutants were checked by PCR and sequencing.

## Acknowledgements

Cryo-EM core: David Bulkey, Glenn Gilbert, Matt Harrington, Eliza N., Li Amber Smith, Yifan Cheng, Feng Wang, David A Agard

## Supplementary Figure Legend

**Supplementary Figure 1: Structural Overview of Stressosome Components.**

(a) RsbS features a conserved Sulfate Transporter and Anti-Sigma factor (STAS) domain, which is integral to its function. (b) The C-terminal domain of RsbR also contains a conserved STAS domain. In addition, its *N*-terminal region houses a non-heme globin domain, which forms turret-like protrusions that extend outward. (c) The STAS domains of RsbR and RsbS interact to form the core architecture of the stressosome. This assembly consists of 20 RsbR dimers and 10 RsbS dimers. The structure shown represents the stressosome from *Listeria monocytogenes*^11^.

**Supplementary Figure 2: Assembly of Stressosome Variants Using Phosphonull Mutants.** Negative-stain electron microscopy reveals particle formation resulting from the interaction of wild-type or mutant RsbR and RsbS proteins with RsbT. The figure includes both a negative stain image and a zoomed-in view of the highlighted region. (a-a1) RsbR_T175A_ST, (b-b1) RsbS_S56A_RT, and (c-c1) RsbR_T209A_ST.

**Supplementary Figure 3: Stressosome Assembly Assessed by Negative-Stain Electron Microscopy of Phosphomimetic Mutants.** (a-c) Negative-stain electron microscopy analysis of particle formation resulting from the interaction of wild-type or phosphomimetic RsbR and RsbS mutants with RsbT. The figure includes both a negative stain image and a zoomed-in view of the highlighted region. Red arrows highlight broken stressosome assemblies or debris. (a-a1) RsbR_T175E_ ST, (b-b1) RsbS_S56D_ RT, and (c-c1) RsbR_T209E_ ST.

**Supplementary Figure 4: Cryo-EM Analysis of Mutant *Listeria monocytogenes* Stressosome Structures.** (a–e) Fourier Shell Correlation (FSC) curves illustrating resolution estimates for cryo-electron microscopy reconstructions of stressosome complexes with the following phosphonull and phosphomimetic mutations: (a) RsbR_T175A_ST, (b) RsbR_T175E_ST, (c) RsbS_S56A_RT, (d) RsbR_T209A_ST, and (e) RsbR_T209E_ST.

**Supplementary Figure 5: Structural Insights into the Native *Listeria* Stressosome Dimer-Dimer Interface (PDB ID: 6QCM)**^11^. Overview of Dimer-Dimer Interactions of the native stressosome assembly: (a) A broad view of the primary trimer of dimers. The detailed views of these interfaces are shown in panels b–d. Dimer-Dimer interfaces are labeled i, ii, and iii. The RsbR and RsbS dimer-dimer interfaces are held together by salt bridges and hydrogen bonds.

