## Supplementary figures and images for "Phosphorylation State Dictates Bacterial Stressosome Assembly and Function"

### Supplementary Figure 1

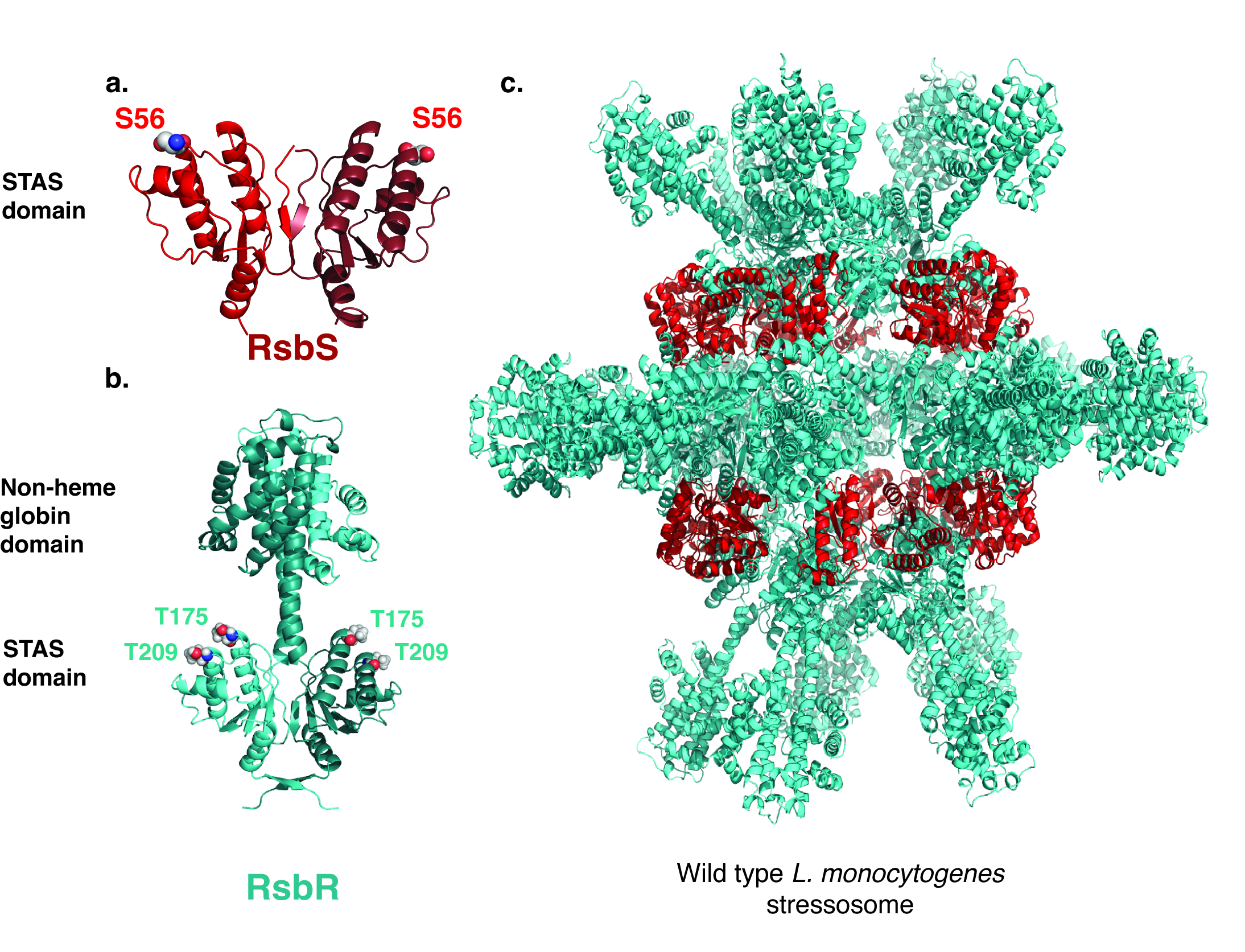

### Supplementary Figure 2

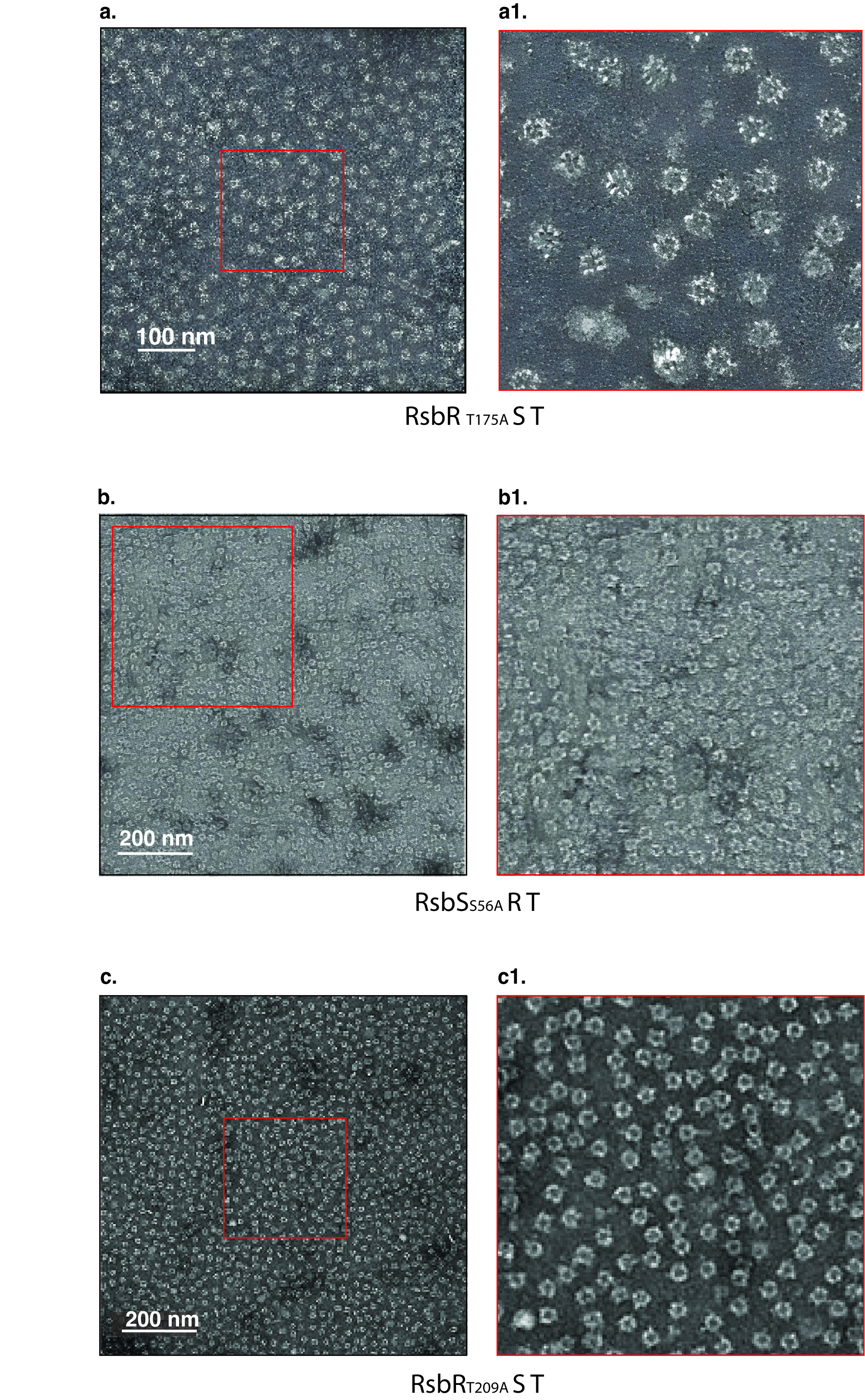

### Supplementary Figure 3

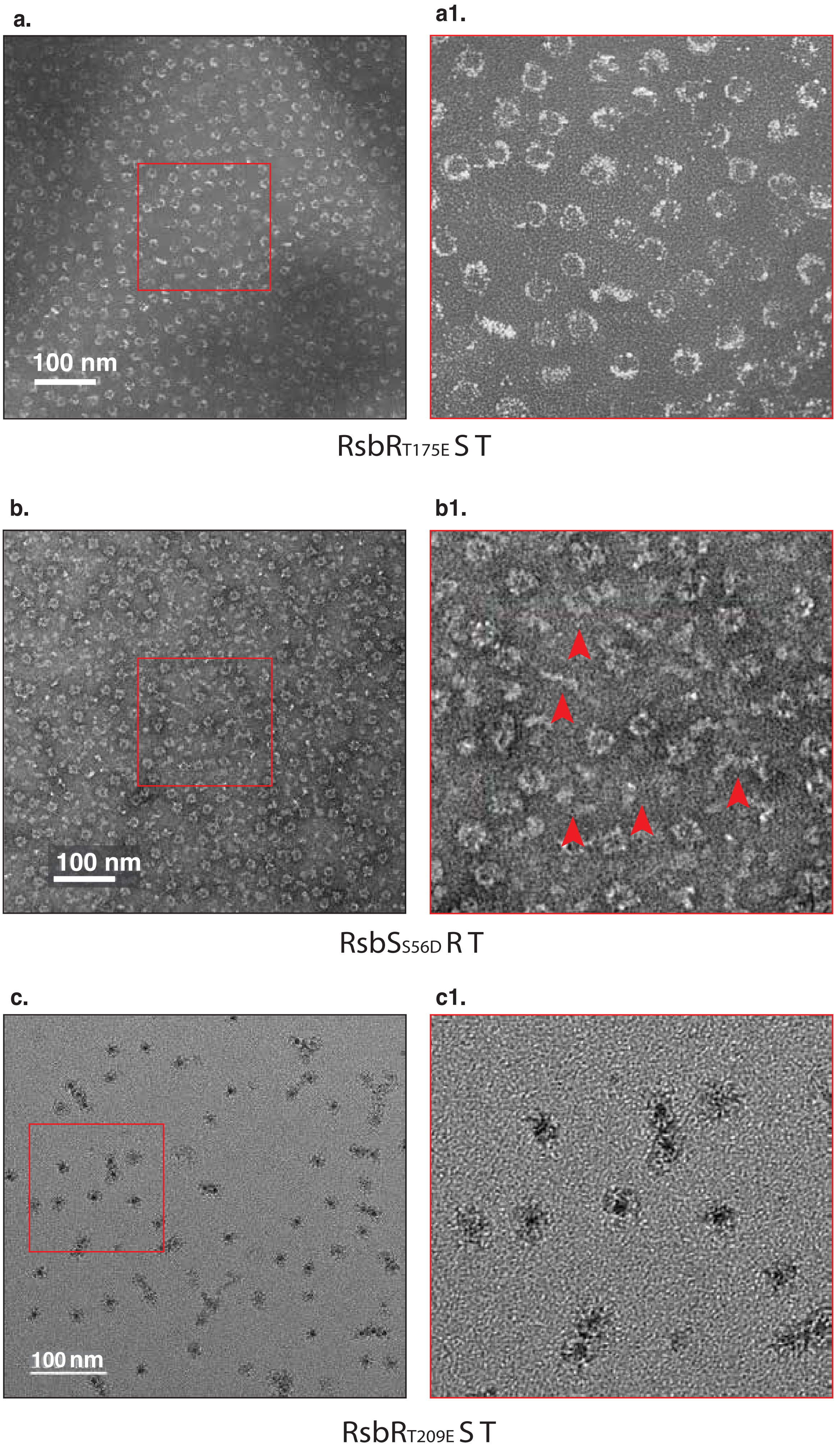

### Supplementary Figure 4

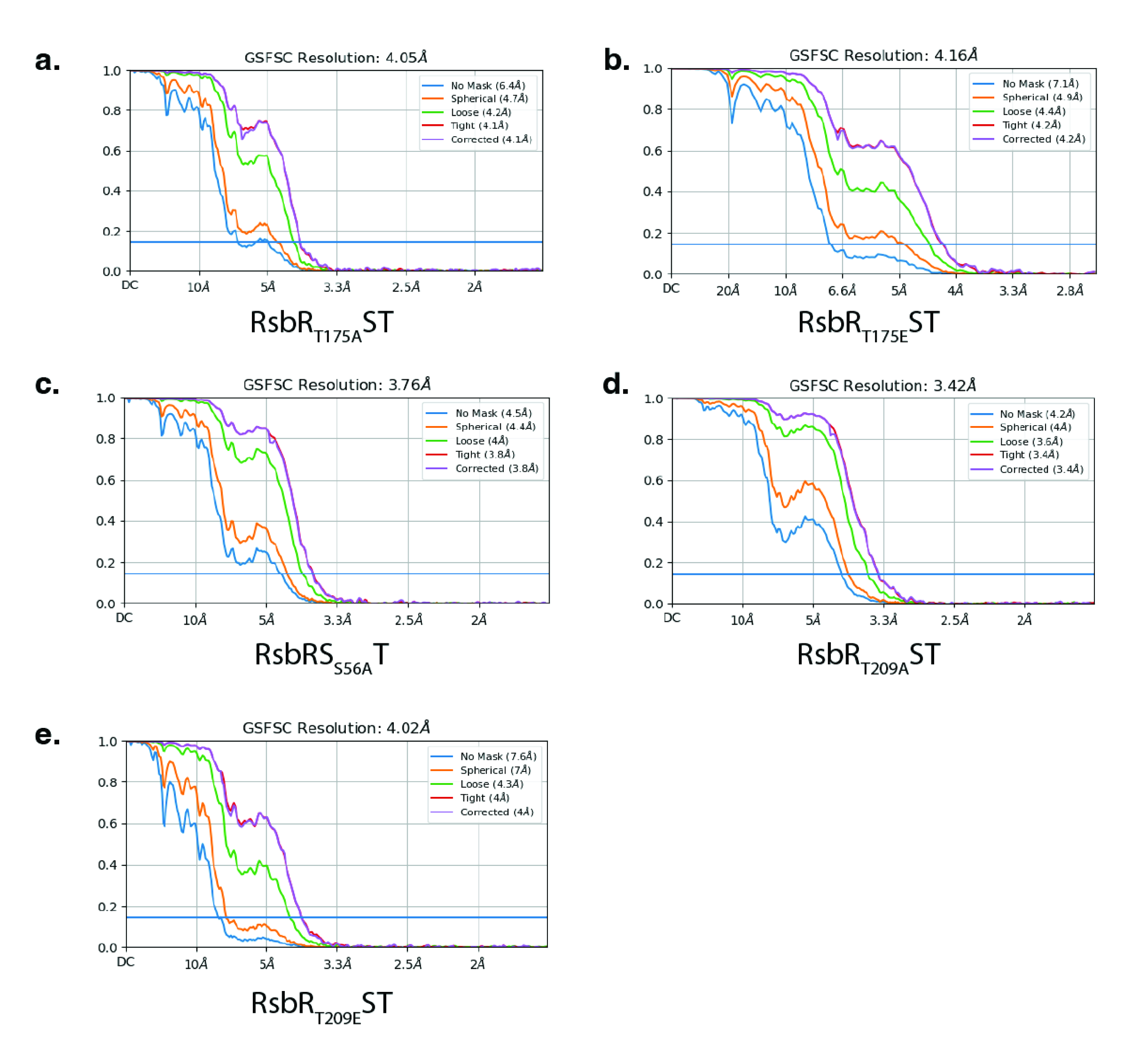

### Supplementary Figure 5

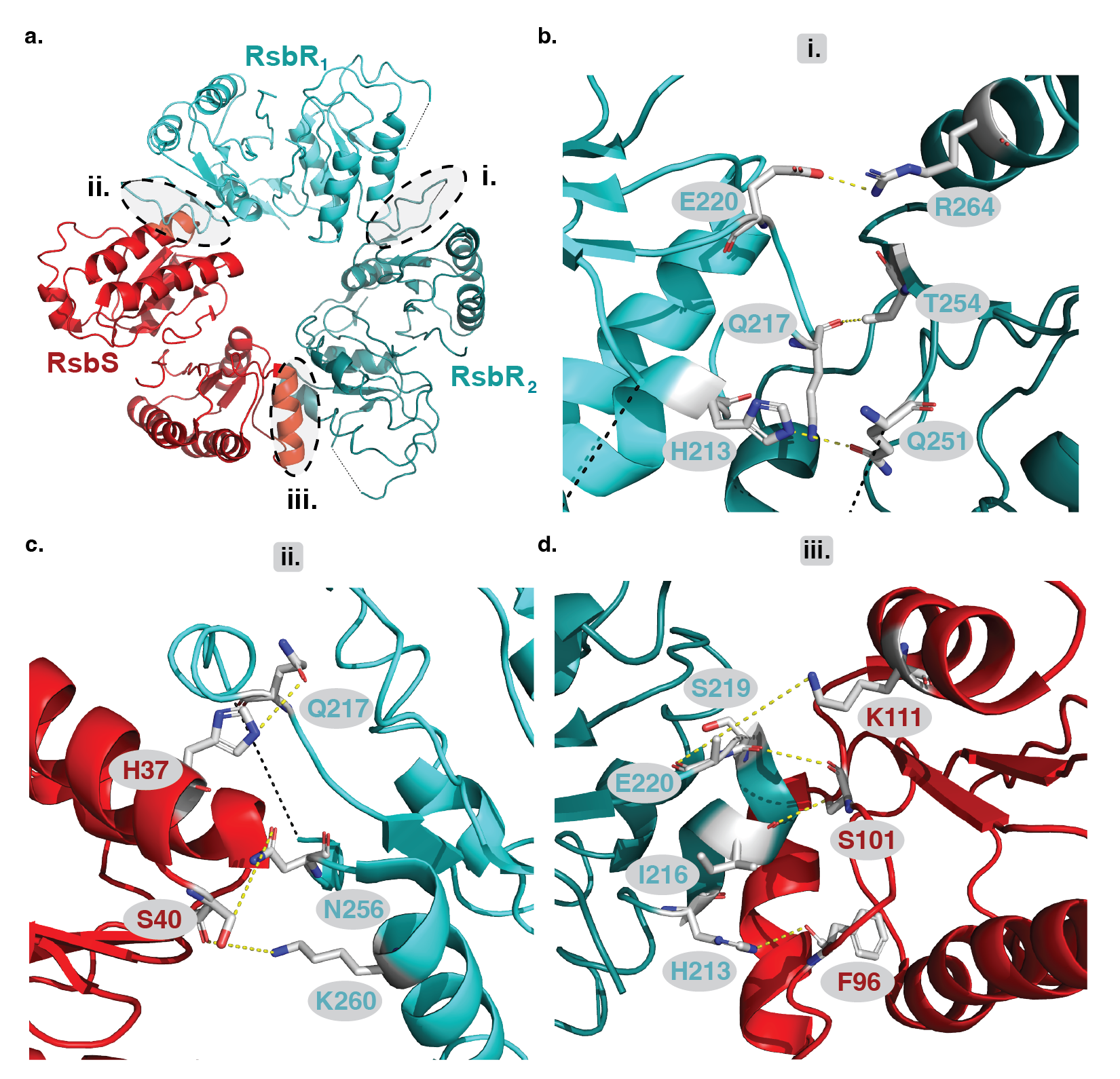
