## Supplementary Table 1 for "Phosphorylation State Dictates Bacterial Stressosome Assembly and Function"

**Supplementary Table 1. Interface Description of Two Units in RsbR$_{T209E}$ST Assembly**

| Interface | Unit 1 (a-d) | | Unit 2 (e-h) | |
|---|---|---|---|---|
| | **Hydrogen Bonding** | **Salt Bridges** | **Hydrogen Bonding** | **Salt Bridges** |
| **Interface i** | $RsbR_1$:E195 :: $RsbR_2$:K260 | $RsbR_1$:R223 :: $RsbR_2$:D250 | $RsbS_2$:K70 :: $RsbS_3$:S101 | No salt bridges |
| | | $RsbR_1$:E195 :: $RsbR_2$:K260 | $RsbS_3$:L71 :: $RsbS_3$:S101 | |
| | | | $RsbS_3$:G73 :: $RsbS_3$:S101 | |
| **Interface ii** | $RsbR_1$:E237 :: $RsbS_1$:H37 | $RsbR_1$:E237 :: $RsbS_1$:H37 | $RsbR_3$:R223 :: $RsbS_2$:S101 | No salt bridges |
| | $RsbR_1$:N256 :: $RsbS_1$:S40 | | $RsbR_3$:L224 :: $RsbS_2$:S101 | |
| **Interface iii** | $RsbS_1$:F96 :: $RsbR_2$:R223 | $RsbR_3$:E248 :: $RsbS_3$:S67 | No salt bridges | $RsbR_3$:E248 :: $RsbS_3$:K70 |
| | $RsbS_1$:S101 :: $RsbR_2$:L224 | | | |
