## Supplementary Table 2 for "Phosphorylation State Dictates Bacterial Stressosome Assembly and Function"

**Supplementary Table 2. List of oligonucleotide primers used in the study**

| Primer | Sequence (5'-3') |
|---|---|
| **Oligonucleotides used for *rsbR-rsbS* cloning in pKSV7** | |
| *rsbR-rsbS*-Up-F | tgcaaacttcatccgtacaaataaag |
| *rsbR-rsbS*-Up-R | tccccacgatttacccctctttttctactac |
| *rsbR-rsbS*-Down-F | gtaagaaaaaggggtgaatcctgqgggatacaatc |
| *rsbR-rsbS*-Down-R | Tttttcagctctccttacag |
| *rsbR-rsbS*-Vector-F | tctgctttattgtaccgatgaaagtttgcaaaaatcttttatacatgcatgccagccgctggac |
| *rsbR-rsbS*-Vector-R | Aaatggttcgtaaggagctgagctgaaaaaatggaaaagcttttgggggaaaagctgcaggagcagtggag |
| **Oligonucleotides to create *Listeria monocytogenes* mutants** | |
| *rsbR1*-T175A-F | acgcagaaagagccaagttaatcatag |
| *rsbR1*-T175A-R | tttctatgattaacttggctctttctgcgtcaatcgttccaattaacggc |
| *rsbR1*- T209A-F | tacgggagttcctgttgttgatgcaatggttgcgcaccatattattc |
| *rsbR1*-T209A-R | accattgcatcaacaacaggaactcc |
| *rsbR1*-T209E-F | ttgcgcaccatattattcaag |
| *rsbR1*-T209E-R | cttcggacgcttgaataatatggtgcgcaaccatttcatcaacaacaggaactccc |
| *rsbS*-S56D-F | attgatataacttctatcgattttattgatgattttattgcaaaaattcttggagatg |
| *rsbS*-S56D-R | atcaataaaatcgatagaagttatatcaatgactactcc |
| *rsbS*-S56E-F | attgatataacttctatcgattttattgatgaatttattgcaaaaattcttggagatg |
| *rsbS*-S56E-R | atcaataaaatcgatagaagttatatcaatgactactcc |
| **Oligonucleotides used for site-directed mutagenesis** | |
| *rsbR1*-T175A-F | aacgattgacgcggaaagagcca |
| *rsbR1*-T175A-R | ccaattaacggcattacag |
| *rsbR1*-T175E-F | aacgattgacgaagaaagagccaag |
| *rsbR1*-T175E-R | ccaattaacggcattacag |
| *rsbR1*- T209A-F | tgttgttgatgcgatggttgcgc |
| *rsbR1*-T209A-R | ggaactcccgtaatatcaatc |
| *rsbR1*-T209E-F | tgtgttgtatgcgatggttgcgc |
| *rsbR1*-T209E-R | ggaactcccgtatatcaatc |
| *rsbS*-S56A-F | ttttattgatgcgttttatgcaaaaattcttg |
| *rsbS*-S56A-R | tcgatagaagttatatcaatgac |
| *rsbS*-S56D-F | ttttattgatgcttttattgcaaaaattcttg |
| *rsbS*-S56D-R | tcgatagaagttatatcaatgac |
