## Supplementary Table 3 for "Phosphorylation State Dictates Bacterial Stressosome Assembly and Function"

**Supplementary Table 3. List of *L. monocytogenes* and *E. coli* Strains Used in the Study**

| Strain and Plasmid | Background | Reference |
|---|---|---|
| **Strains** | | |
| | *E. coli* sm10 | Simon R *et al.,* 1983 |
| WT | *L. monocytogenes* 10403S | Becavin *et al.,* 2014 |
| *DP-L7697 rsbR1* T175A | *L. monocytogenes* 10403S | This study |
| *DP-L7698 rsbR1 T209A* | *L. monocytogenes* 10403S | This study |
| *DP-L7699 rsbR1 T209E* | *L. monocytogenes* 10403S | This study |
| *DP-L7700 rsbS S56D* | *L. monocytogenes* 10403S | This study |
| *DP-L7701 rsbS S56E* | *L. monocytogenes* 10403S | This study |
| *DP-L7624*<br>*WT Δstressosome* | *L. monocytogenes* 10403S | Lobanovska *et al*., 2025 (in preparation) |
| **Plasmids** | | |
| pGEX-4T1-*GST-rsbR* | Expression of N-terminally GST-tagged RsbR wild-type protein | This study |
| pGEX-4T1-*GST-rsbR T175A* | Expression of N-terminally GST-tagged RsbR T175A mutant | This study |
| pGEX-4T1-*GST-rsbR T175E* | Expression of N-terminally GST-tagged RsbR T175E mutant | This study |
| pGEX-4T1-*GST-rsbR T209A* | Expression of N-terminally GST-tagged RsbR T209A mutant | This study |
| pGEX-4T1-*GST-rsbR T209E* | Expression of N-terminally GST-tagged RsbR T209E mutant | This study |
| pGEX-4T1-*GST-rsbS* | Expression of N-terminally GST-tagged RsbS wild-type protein | This study |
| pGEX-4T1-*GST-rsbS S56A* | Expression of N-terminally GST-tagged S56A mutant | This study |
| pGEX-4T1-*GST-rsbS S56D* | Expression of N-terminally GST-tagged S56D mutant | This study |
| pGEX-4T1-*GST-rsbT* | Expression of N-terminally GST-tagged RsbT wild-type protein | This study |
